# Genomic diversity and global distribution of *Saccharomyces eubayanus*, the wild ancestor of hybrid lager-brewing yeasts

**DOI:** 10.1101/709535

**Authors:** Quinn K. Langdon, David Peris, Juan I. Eizaguirre, Dana A. Opulente, Kelly V. Buh, Kayla Sylvester, Martin Jarzyna, María E. Rodríguez, Christian A. Lopes, Diego Libkind, Chris Todd Hittinger

## Abstract

*S. eubayanus*, the wild, cold-tolerant parent of hybrid lager-brewing yeasts, has a complex and understudied natural history. The exploration of this diversity can be used both to develop new brewing applications and to enlighten our understanding of the dynamics of yeast evolution in the wild. Here, we integrate whole genome sequence and phenotypic data of 200 *S. eubayanus* strains, the largest collection to date. *S. eubayanus* has a multilayered population structure, consisting of two major populations that are further structured into six subpopulations. Four of these subpopulations are found exclusively in the Patagonian region of South America; one is found predominantly in Patagonia and sparsely in Oceania and North America; and one is specific to the Holarctic ecozone. *S. eubayanus* is most abundant and genetically diverse in Patagonia, where some locations harbor more genetic diversity than is found outside of South America. All but one subpopulation shows isolation-by-distance, and gene flow between subpopulations is low. However, there are strong signals of ancient and recent outcrossing, including two admixed lineages, one that is sympatric with and one that is mostly isolated from its parental populations. Despite *S. eubayanus*’ extensive genetic diversity, it has relatively little phenotypic diversity, and all subpopulations performed similarly under most conditions tested. Using our extensive biogeographical data, we constructed a robust model that predicted all known and a handful of additional regions of the globe that are climatically suitable for *S. eubayanus*, including Europe. We conclude that this industrially relevant species has rich wild diversity with many factors contributing to its complex distribution and biology.

## Introduction

In microbial population genomics, the interplay of human association and natural variation is still poorly understood. The genus *Saccharomyces* is an optimal model to address these questions for eukaryotic microbes, as it contains both partly human-associated species (i.e. *Saccharomyces cerevisiae*) and mostly wild species (e.g. *Saccharomyces paradoxus)*. These two examples also illustrate the complexity of studying yeast population genomics. Much of *S. cerevisiae* population structure is admixed, and several lineages show signatures of domestication (Liti et al. 2009; Schacherer et al. 2009; Gallone et al. 2016; Gonçalves et al. 2016). In contrast, *S. paradoxus* is almost exclusively found in the wild and has a population structure that is correlated with geography (Leducq et al. 2014; Eberlein et al. 2019). Pure isolates of their more distant relative *Saccharomyces eubayanus* have only ever been isolated from wild environments; yet, hybridizations between *S. cerevisiae* and *S. eubayanus* were key innovations that enabled cold fermentation and lager brewing (Libkind et al. 2011; Gibson and Liti 2015; Hittinger et al. 2018; Baker et al. 2019). Other hybrids with contributions from *S. eubayanus* have been isolated from industrial environments (Almeida et al. 2014; Nguyen and Boekhout 2017), indicating that this species has long been playing a role in shaping many fermented products. This association with both natural and domesticated environments makes *S. eubayanus* an excellent model where both wild diversity and domestication can be investigated.

Since the discovery of *S. eubayanus* in Patagonia (Libkind et al. 2011), this species has received much attention, both for brewing applications and understanding the evolution, ecology, population genomics of the genus *Saccharomyces* (Sampaio 2018). In the years since its discovery, many new globally distributed isolates have been found (Bing et al. 2014; Peris et al. 2014; Rodríguez et al. 2014; Gayevskiy and Goddard 2016; Peris et al. 2016; Eizaguirre et al. 2018). Prior research has suggested that *S. eubayanus* is most abundant and diverse in the Patagonian region of South America, where there are two major populations (Patagonia A/Population A/PA and Patagonia B/Population B/PB) that recent multilocus data suggested are further divided into five subpopulations (PA-1, PA-2, PB-1, PB-2, and PB-3) (Eizaguirre et al. 2018). There are two early-diverging lineages, West China and Sichuan, which were identified through multilocus data (Bing et al. 2014) and whose sequence divergences relative to other strains of *S. eubayanus* are nearly that of currently recognized species boundaries (Peris et al. 2016; Sampaio and Gonçalves 2017; Naseeb et al. 2018). A unique admixed lineage has been found only in North America, which has approximately equal contributions from PA and PB (Peris et al. 2014, 2016). Other isolates from outside Patagonia belong to PB, either the PB-1 subpopulation that is also found in Patagonia (Gayevskiy and Goddard 2016; Peris et al. 2016), or a Holarctic-specific subpopulation that includes isolates from Tibet and from North Carolina, USA (Bing et al. 2014; Peris et al. 2016). This Holarctic subpopulation includes the closest known wild relatives of the *S. eubayanus* subgenomes of lager-brewing yeasts (Bing et al. 2014; Peris et al. 2016).

To explore the geographic distribution, ecological niche, and genomic diversity of this industrially relevant species, here, we present an analysis of whole genome sequencing data for 200 *S. eubayanus* strains. This dataset confirms the previously proposed population structure (Peris et al. 2014, 2016; Eizaguirre et al. 2018) and extends the analysis to fully explore genomic diversity. Even though *S. eubayanus* is genetically diverse and globally distributed, there are not large phenotypic differences between subpopulations. This genomic dataset includes evidence of gene flow and admixture in sympatry, as well as admixture in parapatry or allopatry. While *S. eubayanus* has a well-differentiated population structure, isolation by distance occurs within subpopulations that are found globally, as well as within subpopulations restricted to a handful of locations. Much of the genetic diversity is limited to northern Patagonia, but modeling suggests that there are more geographic areas that are climatically suitable for this species, including Europe. *S. eubayanus* maintains genetic diversity over several dimensions, including multiple high-diversity sympatric populations and a low-diversity widespread invasive lineage. The diversity and dispersal of this eukaryotic microbial species mirror observations in plants and animals, including humans, which shows how biogeographical and evolutionary forces can be shared across organismal sizes, big and small.

## Results

### Global and regional *S. eubayanus* population structure and ecology

To expand on existing data (Libkind et al. 2011; Bing et al. 2014; Peris et al. 2014; Rodríguez et al. 2014; Gayevskiy and Goddard 2016; Peris et al. 2016; Eizaguirre et al. 2018), we sequenced the genomes of 174 additional strains of *S. eubayanus*, bringing our survey to 200 *S. eubayanus* genomes. This large collection provides the most comprehensive dataset to date for *S. eubayanus*. We note that the dataset does not contain West China or Sichuan strains (Bing et al. 2014), which were unavailable for study and may constitute a distinct species or subspecies. These strains were globally distributed (Figure 1A), but the majority of our strains were from South America (172 total, 155 newly sequenced here). The next most abundant continent was North America with 26 strains (19 new to this publication). We also analyzed whole genome sequence data for the single strain from New Zealand (Gayevskiy and Goddard 2016) and the single Tibetan isolate with available whole genome sequence data (Bing et al. 2014; Brouwers et al. 2019). The collection sites in South America span from northern Patagonia to Tierra del Fuego (Figure 1B), while the North American isolates have been sparsely found throughout the continent, including the Canadian province of New Brunswick and the American states of Washington, Wisconsin, North Carolina, and South Carolina (Figure 1C).

**Figure 1.**
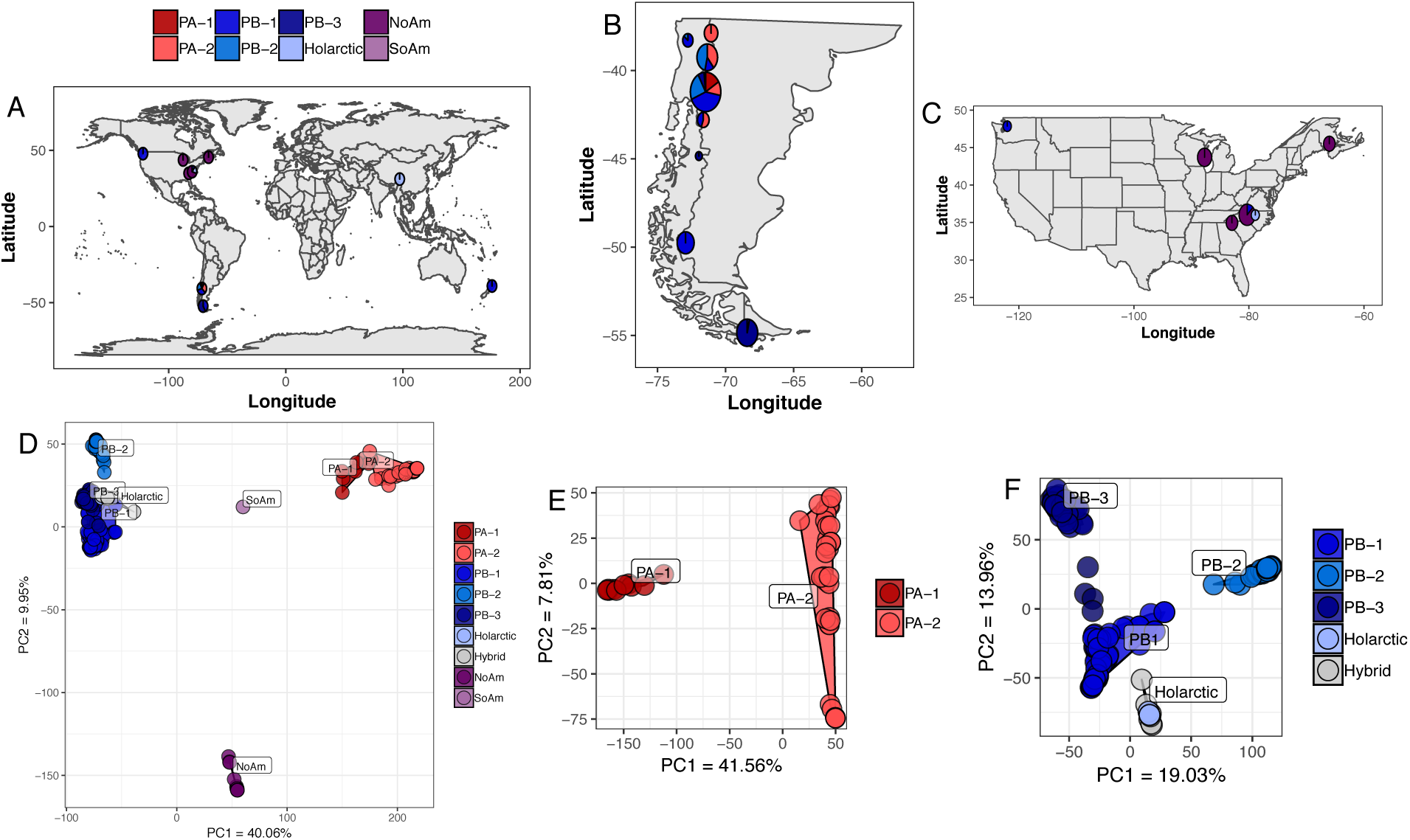
*S. eubayanus* distribution and population structure. *S. eubayanus* has a global distribution and two major populations with six subpopulations. (A) Isolation locations of *S. eubayanus* strains included in the dataset. For visibility, circle size is not scaled by the number of strains. Subpopulation abundance is shown as pie charts. The Patagonian sampling sites have been collapsed to two locations for clarity. Details of sites and subpopulations found in South America (B) and North America (C) with circle size scaled by the number of strains. (D) Whole genome PCA of *S. eubayanus* strains and five hybrids with large contributions from *S. eubayanus*. (E) PCA of just PA. (F) PCA of just PB and hybrid *S. eubayanus* sub-genomes. Color legends in A and D apply to this and all other figures.

To determine population structure, we took several approaches, including Principal Component Analysis (PCA) (Jombart 2008), phylogenomic networks (Huson and Bryant 2006), and STRUCTURE-like analyses (Lawson et al. 2012; Raj et al. 2014). All methods showed that *S. eubayanus* has two large populations that can be further subdivided into a total of six non-admixed subpopulations and one abundant North American admixed lineage (Figure 1D and Figure S1). We previously described the two major populations, PA and PB-Holarctic (Peris et al. 2014, 2016), as well as the subpopulations PA-1, PA-2, PB-1, PB-2, Holarctic, and the North American admixed lineage (Peris et al. 2016). PB-3 had been suggested by multilocus data (Eizaguirre et al. 2018), and our new analyses confirm this subpopulation with whole genome sequence data. All of the strains isolated from outside of South America belonged to either the previously described North American admixed lineage (NoAm) or one of two PB subpopulations, PB-1 or Holarctic. This dataset included novel PB-1 isolates from the states of Washington (yHRMV83) and North Carolina (yHKB35). Unexpectedly, from this same site in North Carolina, we also obtained new isolates of the NoAm admixed lineage (Figure 1C and Table S1), and we obtained additional new NoAm strains in South Carolina. Together, with the North Carolina strains reported here and previously (Peris et al. 2016), this region near the Blue Ridge Mountains harbors three subpopulations or lineages, PB-1, Holarctic, and NoAm. We were also successful in re-isolating the NoAm lineage from the same Wisconsin site, sampling two years later than what was first reported (Peris et al. 2014) (Table S1), indicating that the NoAm admixed lineage is established, not ephemeral, in this location. Additionally, we found one novel South American strain that was admixed between PA (∼45%) and PB (∼55%) (Figure 1D “SoAm”). This global distribution and the well-differentiated population structure of *S. eubayanus* is similar to what has been observed in *S. paradoxus* (Leducq et al. 2014, 2016) and *Saccharomyces uvarum* (Almeida et al. 2014).

*S. eubayanus* has been isolated from numerous substrates and hosts, and our large dataset afforded us the power to analyze host and substrate association by subpopulation. We found that PA-2 was associated with the seeds of *Araucaria araucana* (45.71% of isolates, p-val = 6.11E-07, F-statistic = 15.29). Interestingly, while PB-1 was the most frequently isolated subpopulation (34% of isolates), it has never been isolated from *A. araucana* seeds. Instead, PB-1 was associated with *Nothofagus antarctica* (52.31% of isolates, p-val = 0.017, F-statistic = 3.10). PB-1 was also the subpopulation isolated the most from *Nothofagus dombeyi* (75% of isolates from this tree species), which is a common host of *S. uvarum* (Libkind et al. 2011; Eizaguirre et al. 2018). PB-2 was positively associated with *Nothofagus pumilio* (36.59% of isolates, p-val = 9.60 E-04, F-statistic = 6.59), which could be an ecological factor keeping PB-2 partly isolated from its sympatric subpopulations, PA-2 and PB-1 (Figure 1C). PB-3 was associated with the fungal parasite *Cyttaria darwinii* (14.29% of isolates, p-val = 0.039, F-statistic = 25.34) and *Nothofagus betuloides* (28.57% of isolates, p-val = 5.02E-06, F-statistic = 60.35), which is only found in southern Patagonia and is vicariant with *N. dombeyi*, a host of PB-1. PB-3 was frequently isolated in southern Patagonia (49% of southern isolates) (Eizaguirre et al. 2018), and its association with a southern-distributed tree species could play a role in its geographic range and genetic isolation from the northern subpopulations. Neither *Nothofagus* nor *A. araucana* are native to North America, and we found that our North American isolates were from multiple diverse plant hosts, including *Juniperus virginiana, Diospyros virginiana, Cedrus* sp., and *Pinus* sp. (Table S1), as well as from both soil and bark samples. In Patagonia, *S. eubayanus* has been isolated from exotic *Quercus* trees (Eizaguirre et al. 2018), so even though *Nothofagus* and *A. araucana* are common hosts, *S. eubayanus* can be found on a variety of hosts and substrates. These observed differences in host and substrate could be playing a role in the maintenance of its population structure, especially in sympatric regions of Patagonia.

### All subpopulations grow at freezing temperatures and on diverse carbon sources

*S. eubayanus* comes from a wide range of environments, so we tested if there were phenotypic differences between these subpopulations. We measured growth rates on several carbon sources and stress responses for a large subset of these strains (190) and 26 lager-brewing strains (Figure 2 and Figure S2). Lager-brewing strains grew faster on maltotriose than all subpopulations (p-val < 0.05, Figure 2A), which is consistent with this sugar being one of the most abundant in brewing wort but rare in nature (Salema-Oom et al. 2005). The Holarctic subpopulation grew slower on glucose and maltose compared to all other subpopulations (p-val < 0.05, Figure 2A, Table S2). Overall, the admixed NoAm lineage performed better than PB-1 (p-val = 0.038, Figure 2A), but there was no interaction with carbon source. Therefore, the admixed lineage’s robustness in many conditions could play a role in its success in far-flung North American sites where no pure PA or PB strains have ever been found.

**Figure 2.**
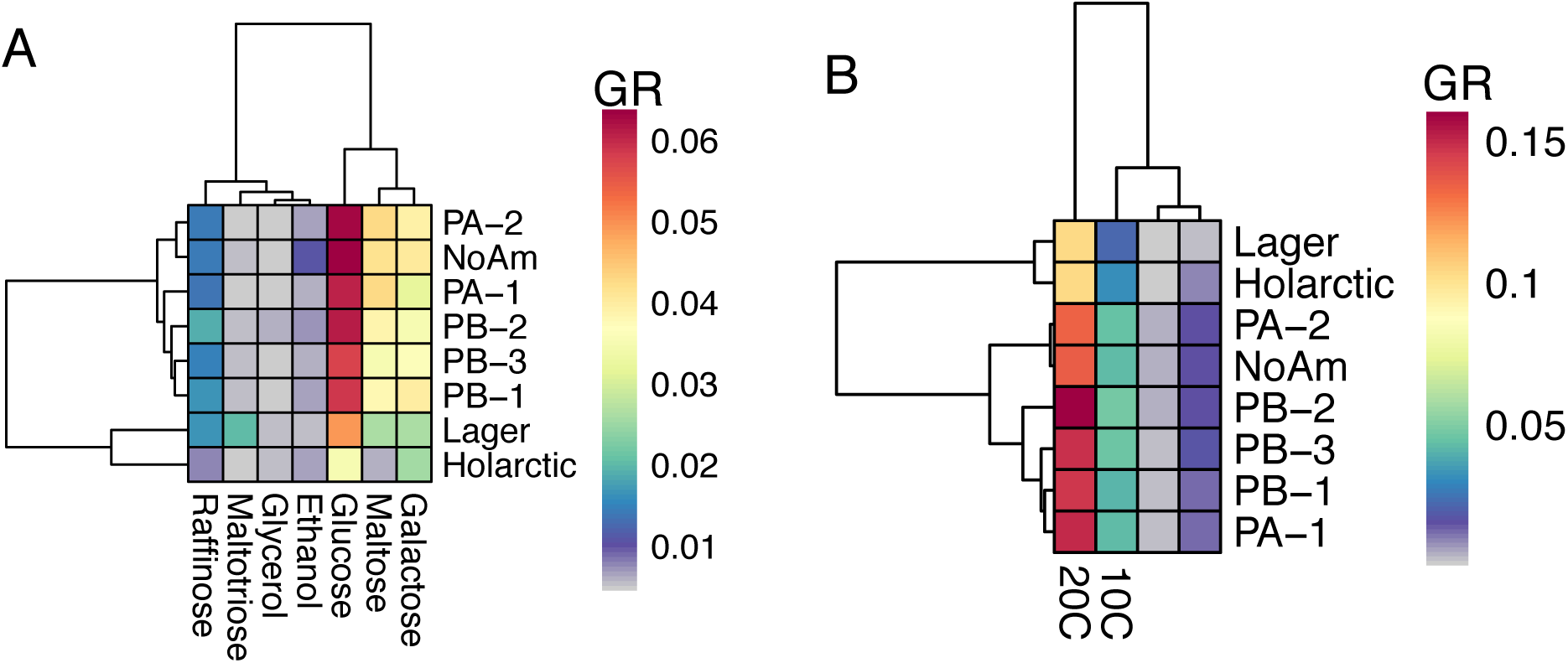
Phenotypic differences. (A) Heat map of mean of maximum growth rate (change in OD/hour) (GR) on different carbon sources by subpopulation. Warmer colors designate faster growth. (B) Heat map of log_10_ normalized growth at different temperatures by subpopulation.

Since *S. eubayanus*’ contribution to the cold-adaptation of hybrid brewing strains is well established (Libkind et al. 2011; Gibson et al. 2013; Baker et al. 2019), we measured growth at 0°C, 4°C, 10°C, and 20°C. All subpopulations grew at temperatures as low as 0°C (Figure 2B and Figure S2), and all *S. eubayanus* subpopulations outperformed lager-brewing yeasts (p < 0.05). Within pure *S. eubayanus*, there were no temperature by subpopulation interactions, indicating that no subpopulation is more cryotolerant than any other subpopulation. In summary, we found that all strains that we tested grew similarly in many environments, and despite the large amount of genotypic diversity observed for this species, we observed much less phenotypic diversity (Figure 2).

### Subpopulations are well differentiated

The mating strategies and life cycle of *Saccharomyces*, with intratetrad mating and haploselfing, often lead to homozygous diploid individuals (Hittinger 2013). Nonetheless, in *S. cerevisiae*, many industrial strains are highly heterozygous (Gallone et al. 2016; Gonçalves et al. 2016; Peter et al. 2018). Here, we analyzed genome-wide heterozygosity in our collection of 200 strains and found only one individual with more than 20,000 heterozygous SNPs (Figure S3). When we phased highly heterozygous regions of its genome and analyzed the two phases separately, we found that both phases grouped within PB-1 (Figure S3C). Thus, while this strain is highly heterozygous, it has contributions from only one subpopulation.

This large collection of strains is a powerful resource to explore natural variation and population demography in a wild microbe, so we analyzed several common population genomic statistics in 50-kbp windows across the genome. We found that diversity was similar between subpopulations (Figure S4A). We also calculated Tajima’s D and found that the genome-wide mean was zero or negative for each subpopulation (Figure S4B), which could be indicative of population expansions. In particular, the most numerous and widespread subpopulation, PB-1, had the most negative and consistent Tajima’s D, suggesting a recent population expansion is especially likely in this case.

For the non-admixed lineages, genome-wide average F_ST_ was consistently high across the genome (Figure S4C). In pairwise comparisons of F_ST,_ PB-1 had the lowest values of any subpopulation (Figure 3A, Figure S4D). These pairwise comparisons also showed that, within each population, there has been some gene flow between subpopulations, even though the subpopulations were generally well differentiated. Linkage disequilibrium (LD) decay indicated low recombination in these wild subpopulations (Figure 3B), with variability between subpopulations. For the species as a whole, LD decayed to one-half at about 5 kbp, which is somewhat higher than the 500bp – 3kbp observed in *S. cerevisiae* (Liti et al. 2009; Schacherer et al. 2009; Peter et al. 2018) and lower than the 9 kbp observed in *S. paradoxus* (Liti et al. 2009), indicating that there is less mating, outcrossing, and/or recombination in this wild species than *S. cerevisiae* and more than in *S. paradoxus*.

**Figure 3.**
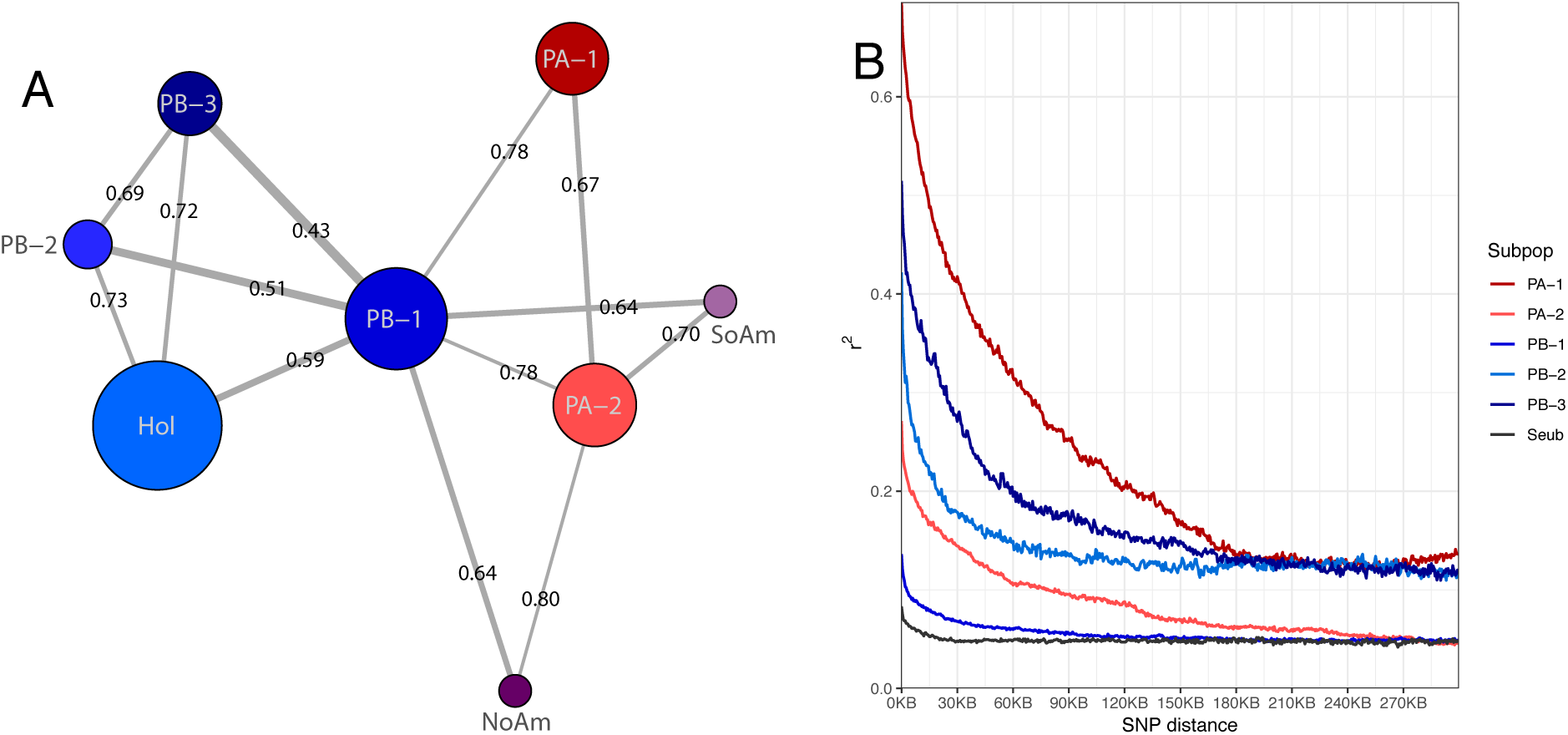
Population genomic parameters. (A) Network built with pairwise F_ST_ values < 0.8 between each subpopulation. F_ST_ values are printed and correspond to line thickness, where lower values are thicker. Circle sizes correspond to genetic diversity. (B) LD decay for each subpopulation (colors) and the species in whole (black).

### Recent admixture and historical gene flow between populations

We previously reported the existence of 7 strains of an admixed lineage in Wisconsin, USA, and New Brunswick, Canada (Peris et al. 2014, 2016). Here, we present 14 additional isolates of this same admixed lineage. These new isolates were from the same site in Wisconsin, as well as two new locations in North Carolina and South Carolina (Table S1). Strikingly, all 21 strains shared the exact same genome-wide ancestry profile (Figure 4A), indicating that they all descended from the same outcrossing event between the two main populations of *S. eubayanus*. These admixed strains were differentiated by 571 SNPs, which also delineated these strains geographically (Figure 4B). Pairwise diversity and F_ST_ comparisons across the genomes suggest that the PA parent came from the PA-2 subpopulation (Figure 4C and Figure S5A) and that the PB parent was from the PB-1 subpopulation (Figure 4D and Figure S5A).

**Figure 4.**
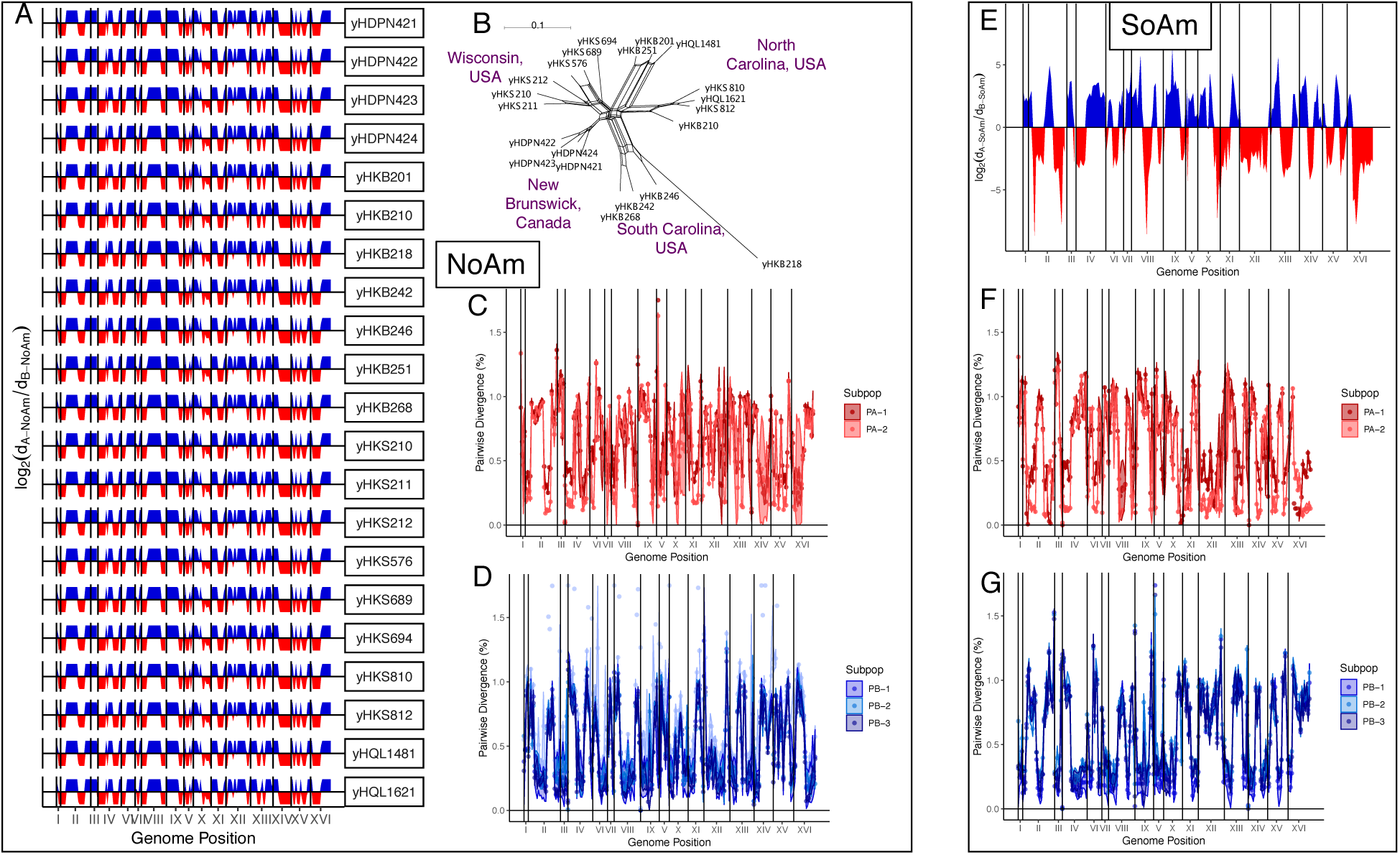
Genomic ancestries of NoAm and SoAm admixed lineages. (A) For all 21 NoAm admixed strains, log_2_ ratio of the minimum PB-NoAm pairwise nucleotide sequence divergence (dB-NoAm) and the minimum PA-NoAm pairwise nucleotide sequence divergence (dA-NoAm) in 50-kbp windows. Colors and log_2_ < 0 or > 0 indicate that part of the genome is more closely related to PA or PB, respectively. (B) Neighbor-Net phylogenetic network reconstructed with the 571 SNPs that differentiate the NoAm strains. The scale bar represents the number of substitutions per site. Collection location is noted. (C) Pairwise nucleotide sequence divergence of the NoAm strain yHKS210 compared to strains from the PA-1 and PA-2 subpopulations of PA in 50-kbp windows. (D) Pairwise nucleotide sequence divergence of the NoAm strain yHKS210 compared to strains from the PB-1, PB-2, and PB-3 subpopulations of PB in 50-kbp windows. (E) log_2_ ratio of the minimum PB-SoAm pairwise nucleotide sequence divergence (dB-SoAm) and the minimum PA-SoAm pairwise nucleotide sequence divergence (dA-SoAm) in 50-kbp windows. Colors and log_2_ < 0 or > 0 indicate that part of the genome is more closely related to PA or PB, respectively. (F) Pairwise nucleotide sequence divergence of the SoAm strain compared to strains from the PA-1 and PA-2 subpopulations of PA in 50-kbp windows. (G) Pairwise nucleotide sequence divergence of the SoAm strain compared to strains of the PB-1, PB-2, and PB-3 subpopulations of PB in 50-kbp windows.

Here, we report a second instance of recent outcrossing between PA and PB. One other strain with fairly equal contributions from the two major populations, PA (∼45%) and PB (∼55%) (Figure 4E), was isolated from the eastern side of Nahuel Huapi National Park, an area that is sympatric for all subpopulations found in South America. This strain had a complex ancestry, where both PA-1 and PA-2 contributed to the PA portions of its genome (Figure 4F and Figure S5B), indicating that its PA parent was already admixed between PA-1 and PA-2. As with the NoAm admixed strains, the PB parent was from the PB-1 subpopulation (Figure 4G and Figure S5B). Together, these two admixed lineages show that outcrossing occurs between the two major populations, and that admixture and gene flow are likely ongoing within sympatric regions of South America.

We also found examples of smaller tracts of admixture between PA and PB that were detectable as 2-12% contributions. These introgressed strains included the taxonomic type strain of *S. eubayanus* (CBS12357^T^), whose genome sequence was mostly inferred to be from PB-1, but it had a ∼4% contribution from PA-1 (Figure S6). We found several other examples of admixture between PA and PB, as well as admixture between subpopulations of PA or of PB (Table S3). Notably, the PB contributions were usually from PB-1, the subpopulation with the largest range, most hosts, and strongest signature of population expansion, factors that would tend to make contact with other subpopulations more likely.

In our collection of 200 strains, we observed nuclear genome contributions from *S. uvarum* in four strains. These four strains all shared the same introgression of ∼150-kbp on chromosome XIV (Figure S7A&B). When we analyzed the portion of the genome contributed by *S. eubayanus*, we found that these strains were all embedded in the PB-1 subpopulation (Figure S7C). Analysis of the 150-kbp region from *S. uvarum* indicated that the closest *S. uvarum* population related to these introgressed strains was SA-B (Figure S7D), a population restricted to South America that has not previously been found to contribute to any known interspecies hybrids (Almeida et al. 2014). These strains thus represent an independent hybridization event between South American lineages of these two sister species that is not related to any known hybridization events among industrial strains (Almeida et al. 2014). These strains show that *S. eubayanus* and *S. uvarum* can and do hybridize in the wild, but the limited number (n=4) of introgressed strains, small introgression size (150-kbp), and shared breakpoints suggest that the persistence of hybrids in the wild is rare.

### Northern Patagonia is a diversity hot spot

Patagonia harbors the most genetic diversity of *S. eubayanus* in our dataset, and four subpopulations were found only there: PA-1, PA-2, PB-2, and PB-3 (Figure 1A and 4A). Therefore, we examined the genetic diversity and range distributions of the isolates from South America more closely. Nahuel Huapi National Park yielded isolates from all five subpopulations found in South America, was the only place where PA-1 was found, and was the location where the SoAm admixed strain was isolated (Figure 5A & B). All five sub-populations were found north of 43°S, an important boundary during the last glaciation period that affects many organisms (Mathiasen and Premoli 2010; Premoli et al. 2010; Quiroga and Premoli 2010). Species-wide, there was more genetic diversity north of this boundary (Figure 5B). In contrast, only PB-1 and PB-3 were found south of 43°S, with both distributions reaching Tierra del Fuego. The southernmost strains were primarily PB-3 (89.7%), but they included two highly admixed PB-1 × PB-3 strains (Table S1 & S3).

**Figure 5.**
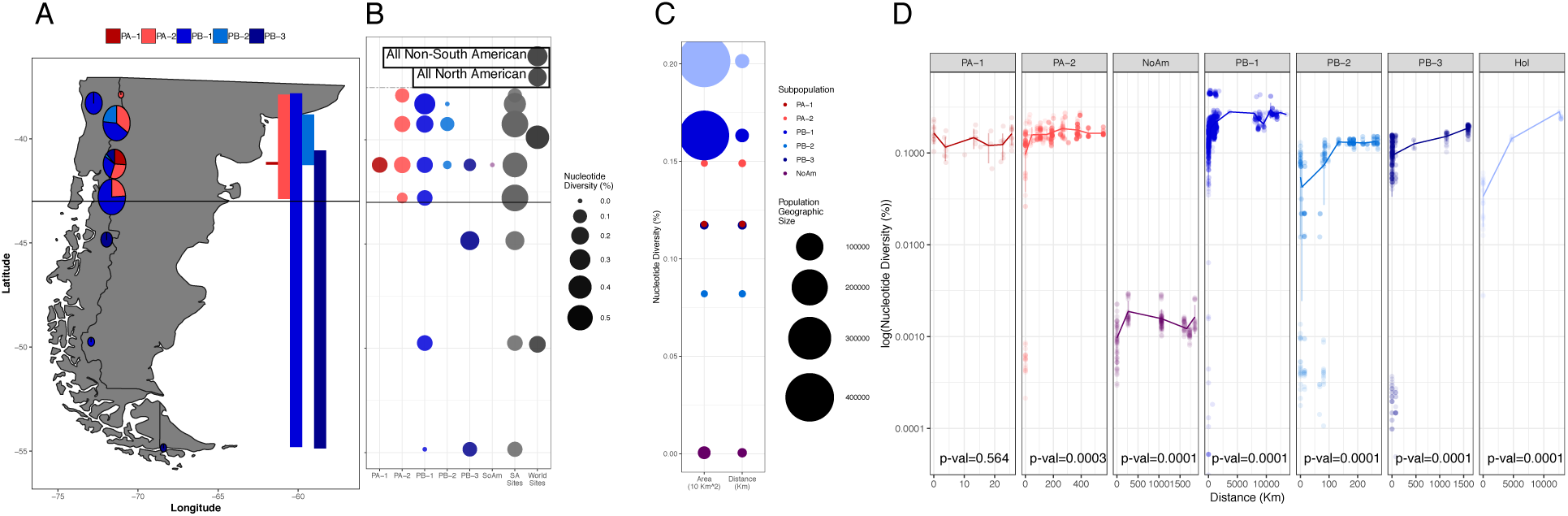
South American genomic diversity versus range, diversity by area, and isolation by distance. (A) Range and genomic diversity of South American sampling sites. Circle sizes correspond to nucleotide diversity of all strains from that site, and pie proportions correspond to each subpopulation’s contribution to *π* at each site. Latitudinal range of each subpopulation is shown to the right. (B) Nucleotide diversity by subpopulation by sampling site, where larger and darker circles indicate more diversity. “SA Sites” in gray show the diversity of all strains found in each South American (SA) site. “World Sites” in darker gray show the nucleotide diversity of all North American or non-South American strains, regardless of subpopulation, compared to South American strains south or north of 43°S, aligned to mean latitude of all strains included in the analysis. (C) Correlation of nucleotide diversity and the area or distance a subpopulation covers. The y-axis shows the nucleotide diversity of each subpopulation, and circle sizes correspond to the geographic sizes of the subpopulations on a log10 scale. Note that PA-1 (dark red) is as diverse as PB-3 (dark blue) but encompasses a smaller area. (D) log_10_(pairwise nucleotide diversity) correlated with distance between strains, which demonstrates isolation by distance. Note that y-axes are all scaled the same but not the x-axes. Holarctic includes the *S. eubayanus* sub-genome of two lager-brewing strains. Figure S8A shows the individual plots for the NoAm lineage. Figure S8B shows the individual plot of PB-1.

Despite the limited geographic range of some subpopulations, their genetic diversity was high, and this diversity often did not scale with the geographic area over which they were found (Figure 5C). The widespread distribution of some subpopulations led us to question if there was isolation by distance within a subpopulation (Figure 5D). We used pairwise measures of diversity and geographic distance between each strain and conducted Mantel tests for each subpopulation. All subpopulations showed significant isolation by distance (Table S4), except PA-1, likely because it had the smallest geographic range (25 km). Even the Mantel test for the least diverse lineage, NoAm, was highly significant (p-val = 0.0001, R^2^= 0.106), indicating that each location has been evolving independently after their recent shared outcrossing and dispersal event. Through these pairwise analyses, we also detected two strains from Cerro Ñielol, Chile, that were unusually genetically divergent from the rest of PB-1 and could potentially be a novel lineage (Figure S8).

### Additional global regions are climatically suitable

The sparse but global distribution of *S. eubayanus* raises questions about whether other areas of the world could be suitable for this species. We used the maxent environmental niche modeling algorithm implemented in Wallace (Kass et al. 2018) to model the global climatic suitability for *S. eubayanus*, using GPS coordinates of all known *S. eubayanus* strains published here and estimates of coordinates for the East Asian isolates (Bing et al. 2014). These niche models were built using the WorldClim Bioclims, which are based on monthly temperature and rainfall measures, reflecting both annual and seasonal trends, as well as extremes, such as the hottest and coldest quarters. How climatic variables affect yeast distributions is an understudied area, and building these models allowed us a novel way to explore climatic suitability.

Using all known locations of isolation (Figure 6), we found that the best model (AIC=2122.4) accurately delineated the known distribution along the Patagonian Andes. In North America, the strains from the Olympic Mountains of Washington state and the Blue Ridge region of North Carolina fell within the predicted areas, and interestingly, these sites had yielded pure PB-1 and Holarctic strains. In contrast, some of the NoAm admixed strains were found in regions that were on the border of suitability in this model (New Brunswick and Wisconsin). In Asia, the model predicts further suitable regions along the Himalayas that are west of known locations.

**Figure 6.**
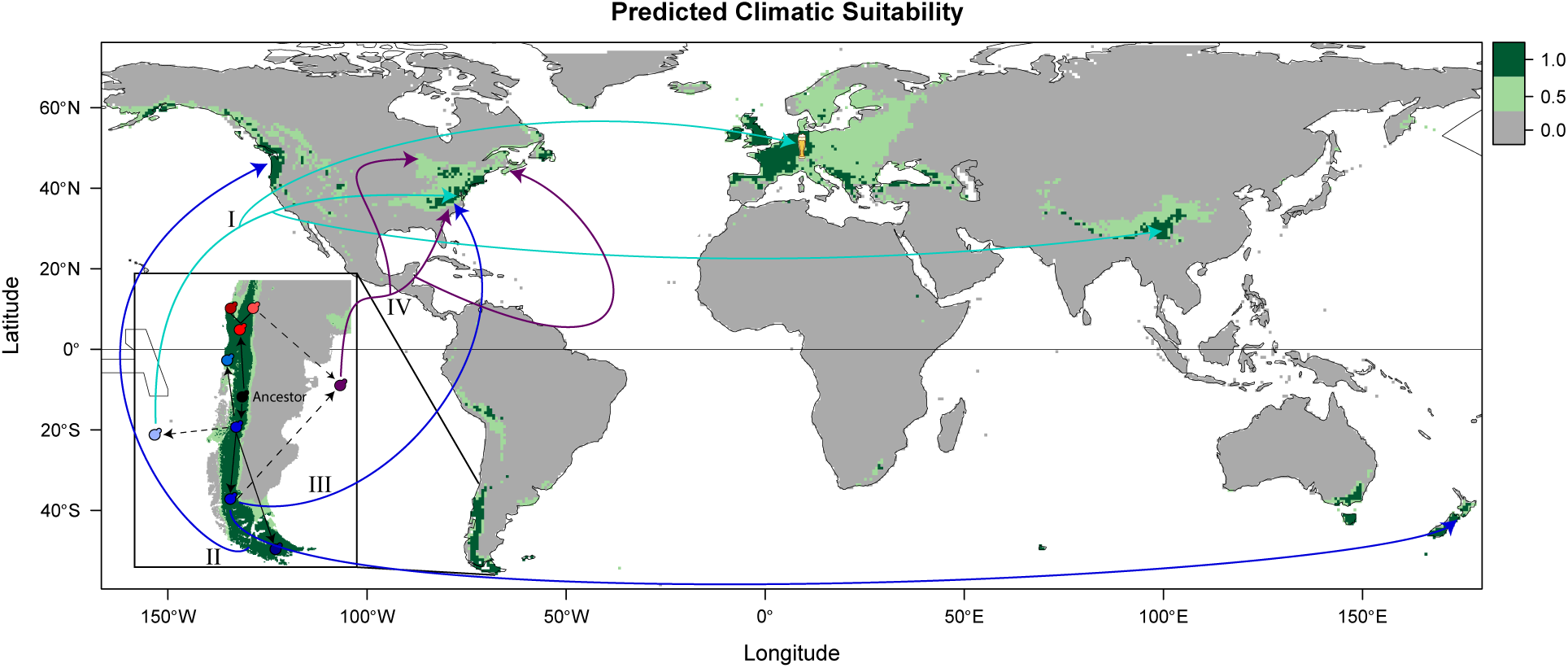
Predicted climatic suitability of *S. eubayanus*. Minimum training presence (light green) and 10^th^ percentile training presence (dark green) based on a model that includes all known *S. eubayanus* isolations, as well as a scenario of dispersal and diversification out of Patagonia (inset and arrows). Black arrows signify diversification events, dotted lines are diversification events where the population is not found in Patagonia, and colored arrows are migration events for the lineage of matching color. Roman numerals order the potential migration events. *S. eubayanus* has not been found in the wild in Europe, but it has contributed to fermentation hybrids, such as lager yeasts. This scenario proposes that the last common ancestor of PA and PB-Holarctic bifurcated into PA (red) and PB-Holarctic (blue), which further radiated into PA-1 (dark red), PA-2 (light red), PB-1 (blue), PB-2 (lighter blue), PB-3 (dark blue), and Holarctic (very light blue). At least four migration events are needed to explain the locations where *S. eubayanus* has been found. I. The Holarctic subpopulation was drawn from the PB-Holarctic gene pool and colonized the Holarctic ecozone. II. PB-1 colonized the Pacific Rim, including New Zealand and Washington state, USA. III. An independent dispersal event brought PB-1 to North Carolina, USA. IV. Outcrossing between PA-2 and PB-1 gave rise to a low-diversity admixed linage that has recently invaded a large swath of North America.

The uneven global distribution of *S. eubayanus* led us to test if models were robust to being built only with the South American locations or only with the non-South American locations (Figure S9). Remarkably, with just the South American isolates, the model (AIC=1327.32) accurately predicted the locations of the non-South American isolates (Figure S9A). Even the model built from the limited number of isolates from outside South America (AIC=558.58) still performed reasonably well, identifying the regions in Patagonia along the Andes where *S. eubayanus* has been found (Figure S9B). Collectively, these models suggest that climatic modeling can predict other suitable regions for eukaryotic microbes. These approaches could be used to direct future sampling efforts or applied to other microbes to gain further insight into microbial ecology.

Notably, all models agree that Europe is climatically a prime location for *S. eubayanus* (Figure S9C), but no pure isolates have ever been found there, only hybrids with *S. uvarum, S. cerevisiae*, or hybrids with even more parents (Almeida et al. 2014). These hybrids with complex ancestries have been found in numerous fermentation environments, suggesting that pure *S. eubayanus* once existed, or still exists at low abundance or in obscure locations, in Europe. Thus, the lack of wild isolates from sampling efforts in Europe remains a complex puzzle.

## Discussion

Here, we integrated genomic, geographic, and phenotypic data for 200 strains of *S. eubayanus*, the largest collection to date, to gain insight into its world-wide distribution, climatic suitability, and population structure. All the strains belong to the two major populations previously described (Peris et al. 2014, 2016), but with the extended dataset, we were able to define considerable additional structure, consisting of six subpopulations and two admixed lineages. These subpopulations have high genetic diversity, high F_ST_, and long LD decay; all measures indicative of large and partly isolated populations undergoing limited gene flow. Despite this high genetic diversity, there was relatively little phenotypic differentiation between subpopulations. This dichotomy between large genetic diversity and limited phenotypic differentiation hints at a complex demographic history where genetically differentiated subpopulations are minimally phenotypically differentiated and grow well in a wide range of environments.

Despite the strong population structure, we also observed considerable evidence of admixture and gene flow. The two recently admixed lineages had nearly equal contributions from the two major populations, but they were the result of independent outcrossing events. The SoAm admixed strain was isolated from a hotspot of diversity and contains contributions from three subpopulations. The NoAm admixed lineage has spread across at least four distant locations, but all strains descended from the same outcrossing event. Since PA has only been isolated in South America, it is intriguing that the NoAm admixed lineage has been successful in so many locations throughout North America. The success of this lineage could be partially explained by its equal or better performance in many environments in comparison to its parental populations (Fig. 2), perhaps contributing to its invasion of several new locations. Several other Patagonian strains also revealed more modest degrees of gene flow between PA and PB. Finally, we characterized a shared nuclear introgression from *S. uvarum* into four Patagonian strains of *S. eubayanus*, demonstrating that hybridization and backcrossing between these sister species has occurred in the wild in South America.

*S. eubayanus* has a paradoxical biogeographical distribution; it is abundant in Patagonia, but it is sparsely found elsewhere with far-flung isolates from North America, Asia, and Oceania. Most subpopulations displayed isolation by distance, but genetic diversity only scaled with geographic range to a limited extent. In Patagonia, some sampling sites harbor more genetic diversity than all non-Patagonian locations together (Figure 5B). Although we found the most genetic diversity and largest number of subpopulations north of 43°S, the pattern of genetic diversity appears to be reversed on the west side of the Andes, at least for the PB-1 subpopulation (Nespolo et al. 2019). This discrepancy could be due to differences in how glacial refugia were distributed (Sérsic et al. 2011) and limitations on gene flow between the east and west sides of the Andes. Together, the levels of *S. eubayanus* genetic diversity found within Patagonia, as well as the restriction of four subpopulations to Patagonia, suggest that Patagonia is the origin of most of the diversity of *S. eubayanus*, likely including the last common ancestor of the PA and PB-Holarctic populations.

The simplest scenario to explain the current distribution and diversity of *S. eubayanus* is a series of radiations in Patagonia, followed by a handful of out-of-Patagonia migration events (Figure 6). Under this model, PA and PB would have bifurcated in Patagonia, possibly in separate glacial refugia. The oldest migration event would have been the dispersal of the ancestor of the Holarctic subpopulation, drawn from the PB gene pool, to the Northern Hemisphere. Multiple more recent migration events could have resulted in the few PB-1 strains found in New Zealand and the USA. The New Zealand and Washington state strains cluster phylogenetically and could have diversified from the same migration event from Patagonia into the Pacific Rim. The PB-1 strain from North Carolina (yHKB35) is genetically more similar to PB-1 strains from Patagonia, suggesting it arrived in the Northern Hemisphere independently of the Pacific Rim strains. Finally, the NoAm admixed strains are likely the descendants of a single, and relatively recent, out-of-Patagonia dispersal. Given that PA appears to be restricted to northern Patagonia, this region could have been where the hybridization leading to the NoAm lineage occurred. While the dispersal vector that brought this admixed lineage to North America is unknown, its far-flung distribution and low diversity show that it has rapidly succeeded by invading new environments.

Other more complex scenarios could conceivably explain the limited number of strains found outside of Patagonia. For example, PA and PB could represent sequential colonizations of Patagonia from the Northern Hemisphere. Under this model, PA would have arrived first and would then have been restricted to northern Patagonia by competition with the later arrival of PB. The Holarctic subpopulation could be interpreted as remnants of the PB population that did not migrate to Patagonia; but the PB-1 strains from the Northern Hemisphere, especially yHKB35, seem far more likely to have been drawn from a Patagonian gene pool than the other way around. Furthermore, the structuring of the PA-1 and the PA-2 subpopulations and of the PB-1, PB-2, and PB-3 subpopulations are particularly challenging to rectify with models that do not allow for diversification within South America. Even more complex scenarios remain possible, and more sampling and isolation will be required to fully elucidate the distribution of this elusive species and more conclusively reject potential biogeographical models.

*S. eubayanus* has a strikingly parallel population structure and genetic diversity to its sister species *S. uvarum* (Almeida et al. 2014; Peris et al. 2016). Both species are abundant and diverse in Patagonia but can be found globally. Both have early diverging lineages, found in Asia or Australasia, that border on being considered novel species. In South America, both have two major populations, where one of these populations is restricted to northern Patagonia (north of 43°S). However, a major difference between the distribution of these species is that pure strains of *S. uvarum* have been found in Europe. Many dimensions of biodiversity could be interacting to bound the distribution and population structure of both *S. eubayanus* and *S. uvarum.* In particular, we know very little about local ecology, including the biotic community and availability of abiotic resources on a microbial scale, but these factors likely all influence microbial success. We show here that substrate and host association vary between subpopulations. In Patagonia, *S. eubayanus* and *S. uvarum* are commonly associated with *Nothofagus*, where *N. dombeyi* is the preferred host of *S. uvarum* (Libkind et al. 2011; Eizaguirre et al. 2018). Therefore, niche partitioning of host trees could be playing role in the persistence of these species in sympatry in Patagonia. However, in locations where *Nothofagus* is not found and there are perhaps fewer hosts, competitive exclusion between the sister species *S. eubayanus* and *S. uvarum*, could influence distribution. Competition for a narrower set of hosts could potentially explain why only *S. uvarum* has been found in Europe as pure strains, while *S. eubayanus* has not. A second factor influencing distribution and population structure could be dispersal. Yeasts could migrate via many avenues, such as wind, insect, bird, or other animals (Francesca et al. 2012, 2014; Stefanini et al. 2012; Gillespie et al. 2012). Human mediated-dispersal has been inferred for the *S. cerevisiae* Wine and Beer lineages and for the *S. paradoxus* European/SpA lineage (Gallone et al. 2016; Gonçalves et al. 2016; Leducq et al. 2014; Kuehne et al. 2007). A third bounding factor could be a region’s historical climate. Glacial refugia act as reservoirs of isolated genetic diversity that allow expansion into new areas after glacial retreat (Stewart and Lister 2001). 43°S is a significant geographic boundary due to past geological and climatic variables (Mathiasen and Premoli 2010; Eizaguirre et al. 2018), and many other species and genera show a distinction between their northern and southern counterparts, including *Nothofagus* (Mathiasen and Premoli 2010; Premoli et al. 2012). *S. eubayanus* and *S. uvarum* diversities are also strongly affected by the 43°S boundary (Almeida et al. 2014; Eizaguirre et al. 2018), and it seems likely that the microbes experienced some of the same glaciation effects as their hosts. The strong correlation of *S. eubayanus* and *S. uvarum* population structures with 43°S further implies a longstanding and intimate association with Patagonia.

The sparse global distribution and complex patterns of genetic diversity continue to raise questions about the niche and potential range of *S. eubayanus*. Our climatic modeling suggests that parts of Europe would be ideal for *S. eubayanus*. Despite extensive sampling efforts, *S. eubayanus* has never been isolated in Europe (Sampaio 2018). However, recent environmental sequencing of the fungal specific ITS1 region hinted that *S. eubayanus* may exist in the wild in Europe (Alsammar et al. 2019). Considerable caution is warranted in interpreting this result because the rDNA locus quickly fixes to one parent’s allele in interspecies hybrids, there is only a single ITS1 SNP between *S. uvarum* and *S. eubayanus*, and the dataset contained very few reads that mapped to *S. eubayanus*. Still, the prevalence of hybrids with contributions from the Holarctic lineage of *S. eubayanus* found in Europe (Peris et al. 2016) suggests that the Holarctic lineage exists in Europe, or at least existed historically, allowing it to contribute to many independent hybridization events.

The patterns of radiation and dispersal observed here mirror the dynamics of evolution found in other organisms (Czekanski-Moir and Rundell 2019), including humans (Nielsen et al. 2017). *S. eubayanus* and humans harbor diverse and structured populations in sub-Saharan Africa and Patagonia, respectively. In these endemic regions, both species show signals of ancient and recent admixture between these structured populations. Both species have successfully colonized wide swaths of the globe, with the consequence of repeated bottlenecks in genetic diversity. While anatomically modern humans underwent a single major out-of-Africa migration that led to the peopling of the world (Nielsen et al. 2017), *S. eubayanus* has experienced several migration events from different populations that have led to more punctate global distribution. For both species, intraspecific admixture and interspecific hybridization appear to have played adaptive roles in the success of colonizing these new locations. In humans, introgressions from past hybridizations with both Neanderthals and Denisovans underlie many adaptive traits (Racimo et al. 2015), while the cold fermentation of lager-brewing would not be possible without the cryotolerance of *S. eubayanus* and the aggressive fermentation of domesticated ale strains of *S. cerevisiae* (Gibson and Liti 2015). These parallels illustrate how the biogeographical and evolutionary dynamics observed in plants and animals also shape microbial diversity. As yeast ecology and population genomics (Marsit et al. 2017; Yurkov 2017) move beyond the Baas-Becking “Everything is everywhere” hypothesis of microbial ecology (Baas-Becking 1934; de Wit and Bouvier 2006), the rich dynamics of natural diversity that is hidden in the soil at our feet is being uncovered.

## Methods

### Wild strain isolations

All South American isolates were sampled, isolated, and identified as described previously (Libkind et al. 2011; Eizaguirre et al. 2018). North American isolates new to this publication were from soil or bark samples from the American states of Washington, Wisconsin, North Carolina, and South Carolina (Table S1). Strain enrichment and isolation was done as previously described (Sylvester et al. 2015; Peris et al. 2016, 2014), with a few exceptions in temperature and carbon source of isolation (Table S1). Specifically, two strains were isolated at 4°C, eight strains were isolated at room temperature, and six strains were isolated on a non-glucose carbon source: three in galactose, two in sucrose, and one in maltose (Table S1).

### Whole genome sequencing and SNP-calling

Whole genome sequencing was completed with Illumina paired-end reads as described previously (Peris et al. 2016; Shen et al. 2018). Reads were aligned to the reference genome (Baker et al. 2015), SNPs were called, masked for low coverage, and retained for downstream analysis as described previously (Peris et al. 2016). One strain, yHCT75, had more than 20,000 heterozygous SNPs called. This strain was pseudo-phased using read-backed phasing in GATK (McKenna et al. 2010) and split into two phases. Short-read data is deposited in the NCBI Short Read Archive under PRJNA555221.

### Population genomic analyses

Population structure was defined using several approaches: `fastSTRUCTURE` (Raj et al. 2014), `fineSTRUCTURE` (Lawson et al. 2012), `SplitsTree v4` (Huson and Bryant 2006), and Principal Component Analysis with the *adegenet* package in `R` (Jombart 2008). `fineSTRUCTURE` analysis was completed using all strains and 11994 SNPs. The `SplitsTree` network was built with this same set of strains and SNPs. `fastStructure` analysis was completed with as subsample of 5 NoAm strains and 150165 SNPs. We tested K=1 through K=10 and selected K=6 using the “`chooseK.py`” command in `fastSTRUCTURE`. All calculations of pairwise divergence, F_ST_, and Tajima’s D for subpopulations were computed using the `R` package *PopGenome* (Pfeifer and Wittelsbuerger 2015) in windows of 50-kbp. Pairwise divergence between strains was calculated across the whole genome using *PopGenome*. LD was calculated using `PopLDdecay` (Zhang et al. 2019). Geographic area and distance of subpopulations was calculated using the *geosphere* package in `R` (Hijmans et al. 2019). The Mantel tests were completed using *ade4* package of `R` (Dray and Dufour 2007). The F_ST_ network was built with *iGraph* in `R` (Csardi and Nepusz 2006).

### Niche projection with Wallace

Climatic modeling of *S. eubayanus* was completed using the `R` package *Wallace* (Kass et al. 2018). Three sets of occurrence data were tested: one that included only GPS coordinates for strains from South America, one that included only non-South American isolates, and one that included all known isolates (Table S1). We could use exact GPS coordinates for most strains, except for the strains from East Asia, where we estimated the locations (Bing et al. 2014). WorldClim bioclimatic variables were obtained at a resolution of 2.5 arcmin. The background extent was set to “Minimum convex polygon” with a 0.5-degree buffer distance and 10,000 background points were sampled. We used block spatial partitioning. The model was built using the Maxent algorithm, using the feature classes: L (linear), LQ (linear quadradic), H (Hinge), LQH, and LQHP (Linear Quadradic Hinge Product) with 1-3 regularization multipliers and the multiplier step value set to 1. The model was chosen based on the Akaike Information Criterion (AIC) score (Table S5). The best models were then projected to the all continents, except Antarctica.

### Phenotyping

Strains were first revived in Yeast Peptone Dextrose (YPD) and grown for 3 days at room temperature. These saturated cultures were then transferred to two 96-well microtiter plates, for growth rate and stress tolerance phenotyping. These plates were incubated overnight. Cells were pinned from these plates into plates for growth rate measurements. For temperature growth assays, cells were pinned into four fresh YPD microtiter plates and then incubated at 0°C, 4°C, 10°C, and 20°C. For the microtiter plates at 0°C, 4°C, and 10°C, OD was measured at least once a day for two weeks or until a majority of the strains had reached stationary phase. Growth on different carbon sources was measured at 20°C in MM media with 2% of the respective carbon source. Carbon sources tested were: glucose, galactose, raffinose, maltose, maltotriose, ethanol, and glycerol. OD was read every two hours for one week or until saturation. All phenotyping was completed in biological triplicate. The carbon source data was truncated to 125 hours to remove artifacts due to evaporation. Growth curves were analyzed using the package *grofit* (Kahm et al. 2010) in `R` to measure saturation and growth rate. We then averaged each strain over the triplicates. We used an ANOVA corrected with Tukey’s HSD to test for growth rate interactions between subpopulation and carbon source or subpopulation and temperature. We used the `R` package *pvclust* (Suzuki and Shimodaira 2006) to cluster and build heatmaps of growth rate by subpopulation.

Heat shock was completed by pelleting 200μl saturated culture, removing supernatant, resuspending in 200μl YPD pre-heated to 37°C, and incubating for one hour at 37°C, with a room temperature control. Freeze-thaw tolerance was tested by placing saturated YPD cultures in a dry ice ethanol bath for two hours, with a control that was incubated on ice. After stress, the strains were serially diluted 1:10 three times and pinned onto solid YPD. These dilution plates were then photographed after 6 and 18 hours. `CellProfiler` (Lamprecht et al. 2007) was used to calculate the colony sizes after 18 hours, and the 3^rd^ (1:1000) dilutions were used for downstream analyses. The heat shock measurements were normalized by the room temperature controls, and the freeze thaw measurements were normalized by the ice incubation controls. Statistical interactions of subpopulations and stress responses were tested as above.

## Data Availability

All short-read genome sequencing data has been deposited in the NCBI Short Read Archive under the PRJNA555221. Accessions of public data is given in Table S1.

## Acknowledgements

We thank Francisco A. Cubillos for coordinating publication with their study; Sean D. <u>Schoville</u>, José Paulo Sampaio, Paula Gonçalves, and members of the Hittinger Lab, in particular EmilyClare P. Baker, for helpful discussion and feedback; Amanda B. Hulfachor and Martin Bontrager for preparing a subset of Illumina libraries; the University of Wisconsin Biotechnology Center DNA Sequencing Facility for providing Illumina sequencing facilities and services; the Agricultural Research Service (ARS) NRRL collection, Christian R. Landry, Ashley Kinart, and Drew T. Doering for strains included in phenotyping; Huu-Vang Nguyen for strains used for hybrid genome comparison; and Leslie Shown, Anita R. & S. Todd Hittinger, EmilyClare P. Baker, and Ryan V. Moriarty for collecting samples and/or isolating strains. This material is based upon work supported by the National Science Foundation under Grant Nos. DEB-1253634 (to CTH) and DGE-1256259 (Graduate Research Fellowship to QKL), the USDA National Institute of Food and Agriculture Hatch Project No. 1003258 to CTH, and in part by the DOE Great Lakes Bioenergy Research Center (DOE BER Office of Science Nos. DE-SC0018409 and DE-FC02-07ER64494 to Timothy J. Donohue). QKL was also supported by the Predoctoral Training Program in Genetics, funded by the National Institutes of Health (5T32GM007133). DP is a Marie Sklodowska-Curie fellow of the European Union’s Horizon 2020 research and innovation program (Grant Agreement No. 747775). DL was supported by CONICET (PIP11220130100392CO), FONCyT (PICT 3677, PICT 2542), Universidad Nacional del Comahue (B199), and NSF-CONICET grant. CTH is a Pew Scholar in the Biomedical Sciences and H. I. Romnes Faculty Fellow, supported by the Pew Charitable Trusts and Office of the Vice Chancellor for Research and Graduate Education with funding from the Wisconsin Alumni Research Foundation (WARF), respectively.

## Author Contributions

QKL, DL, and CTH, conceived of the study; QLK, DP, JIE, DL, and CTH refined concept and design; QKL performed all population genomic and ecological niche analyses; QKL and DAO sequenced genomes, performed phenotyping and statistical analyses, and mentored KVB and MJ; DL and CTH supervised the study; KVB, KS, and MJ isolated and/or identified North American strains; JIE, MER, CAL, and DL isolated and/or provided South American strains; and QKL and CTH wrote the manuscript with editorial input from all co-authors.

---

**Supplementary Figure 1.**
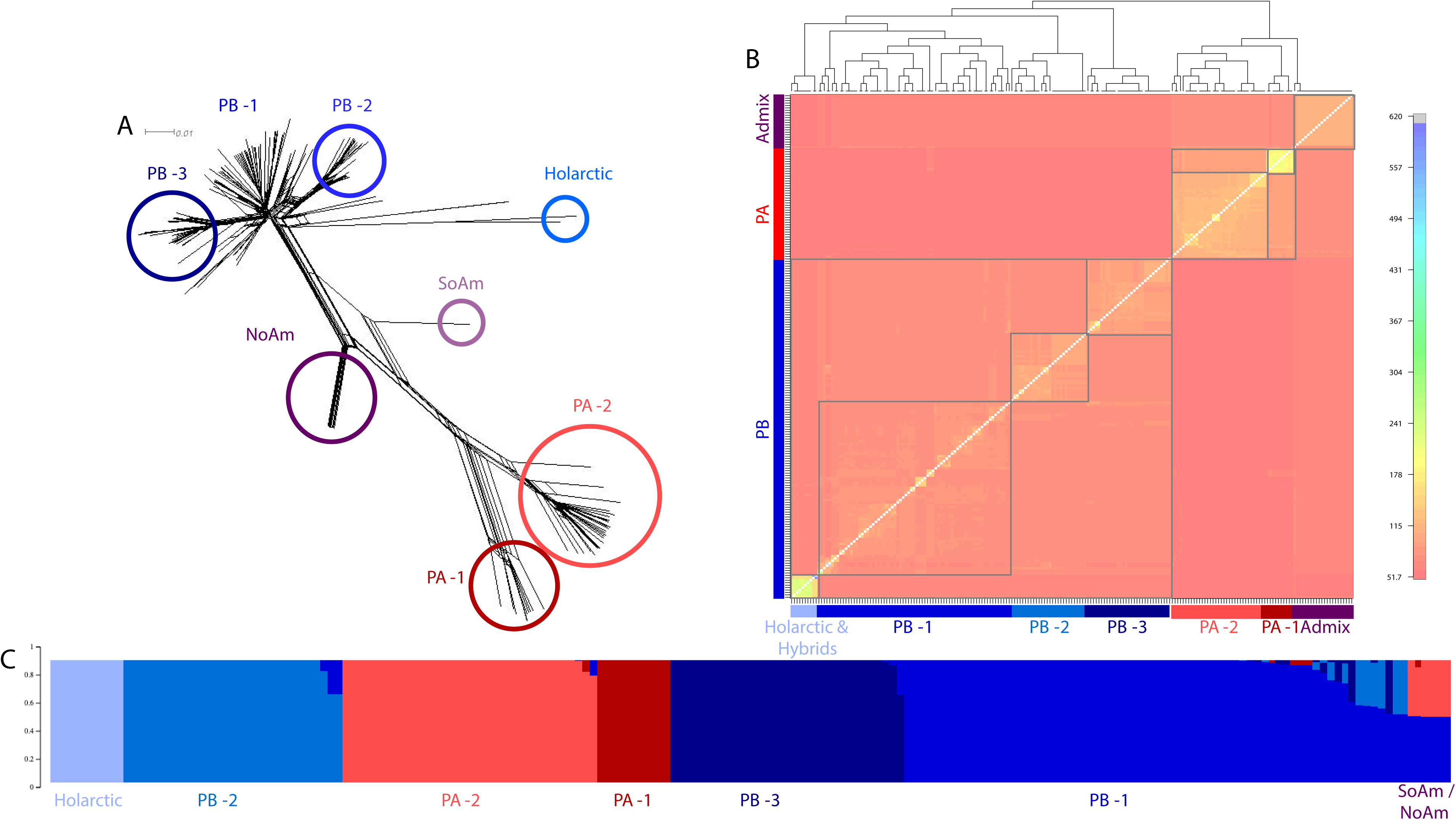
Additional visualizations of population structure. (A) SplitsTree network tree built with 11994 SNPs with subpopulations circled and labeled. (B) FineStructure co-ancestry plot built with 11994 SNPs. Bluer colors correspond to more genetic similarity. Boxes have been added to label the subpopulations. (C) FastSTRUCTURE plot (K=6) built with 150165 SNPs and showing the same six monophyletic subpopulations found with other approaches. Only five NoAm strains were included in the fastSTRUCTURE analysis.

**Supplementary Figure. 2.**
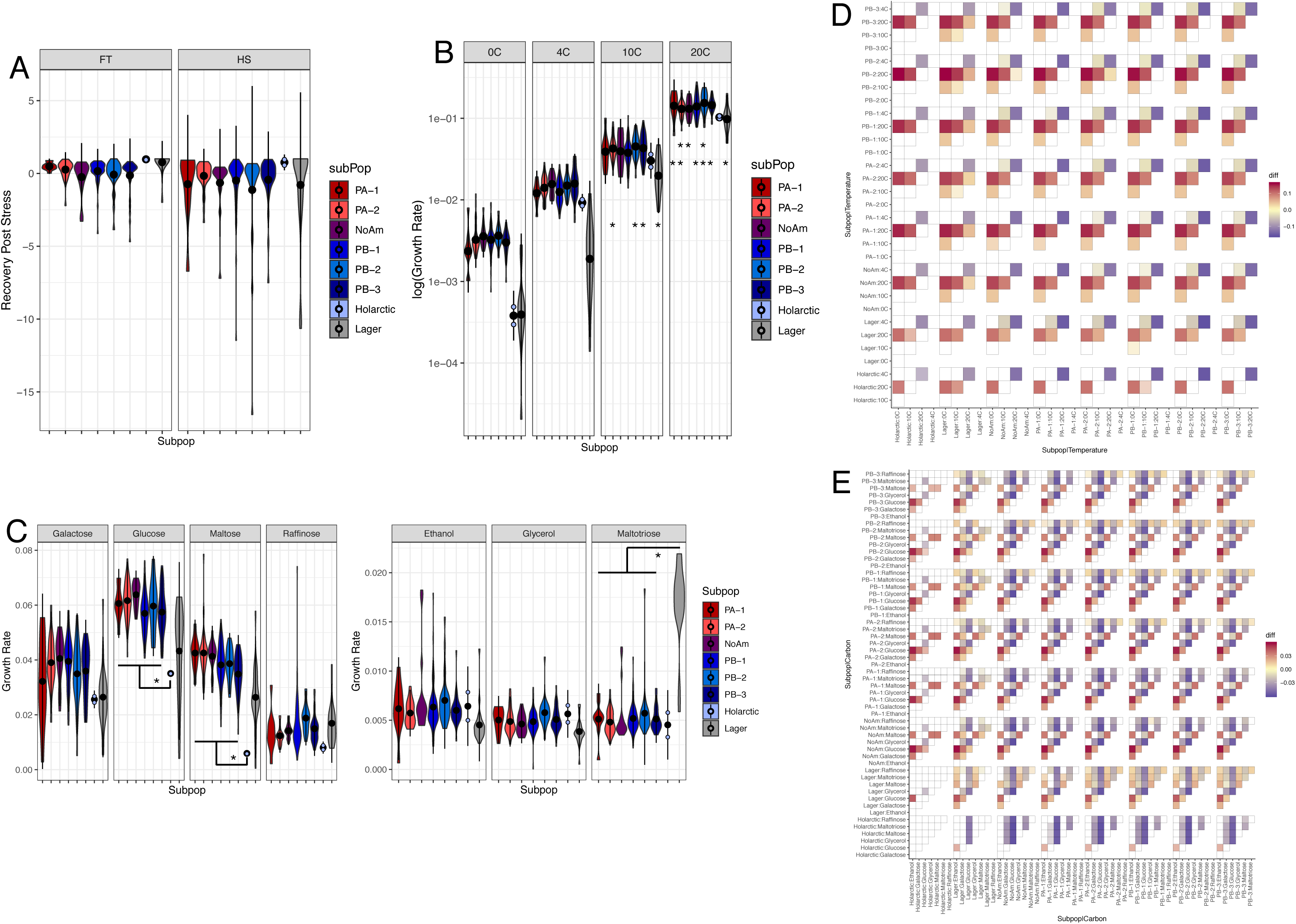
Additional phenotypic data. (A) Violin plots of recovery from stress, normalized by controls. There were no significant subpopulation by stress interactions. (B) Violin plots of log_10_ normalized mean growth rates of each subpopulation at 0°C, 4°C, 10°C, and 20°C. * = p-val < 0.05 of interactions between Lager and PA-2, PB-2, and PB-3 at 10°C; Lager and PA-1, PA-2, PB-1, PB-2, and PB-3 at 20°C; and PB-2 and both PA-2 and NoAm at 20°C (C) Violin plots of mean growth rate on different carbon sources (* = p-val < 0.05). (D) Heatmaps of significant subpopulation by temperature interactions and (E) significant subpopulation by carbon source interactions. Warmer colors indicate that the subpopulation-temperature or carbon source on the left hand had a faster growth rate than the subpopulation-temperature or carbon source along the bottom; cooler colors represent the reverse. Non-significant interactions, based on multiple test corrections, are in white. More intense colors represent smaller p-values.

**Supplementary Figure 3.**
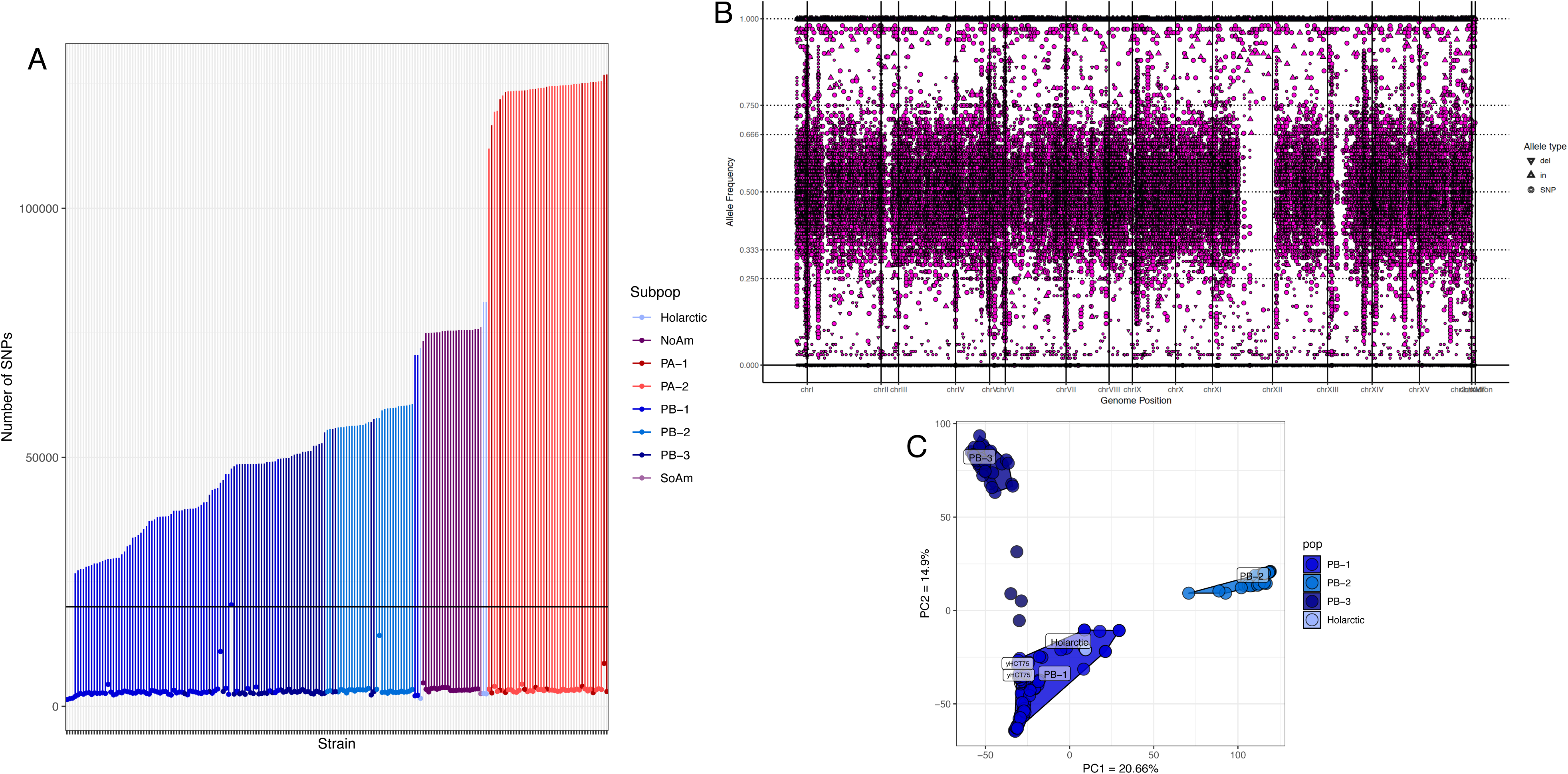
Heterozygosity analyses. (A) Summary of all SNPs vs SNPs called as heterozygous compared to the taxonomic type strain for all 200 strains included in this study. The upper limit of the bar is the total SNP count. The lower point corresponds to SNPs called as heterozygous. The horizontal line is 20k SNPs. (B) Strain yHCT75 (CRUB 1946) is the only strain with > 20K heterozygous SNPs. (C) When the heterozygous SNPs of yHCT75 were pseudo-phased (labeled), both phases clustered with PB-1.

**Supplementary Figure 4.**
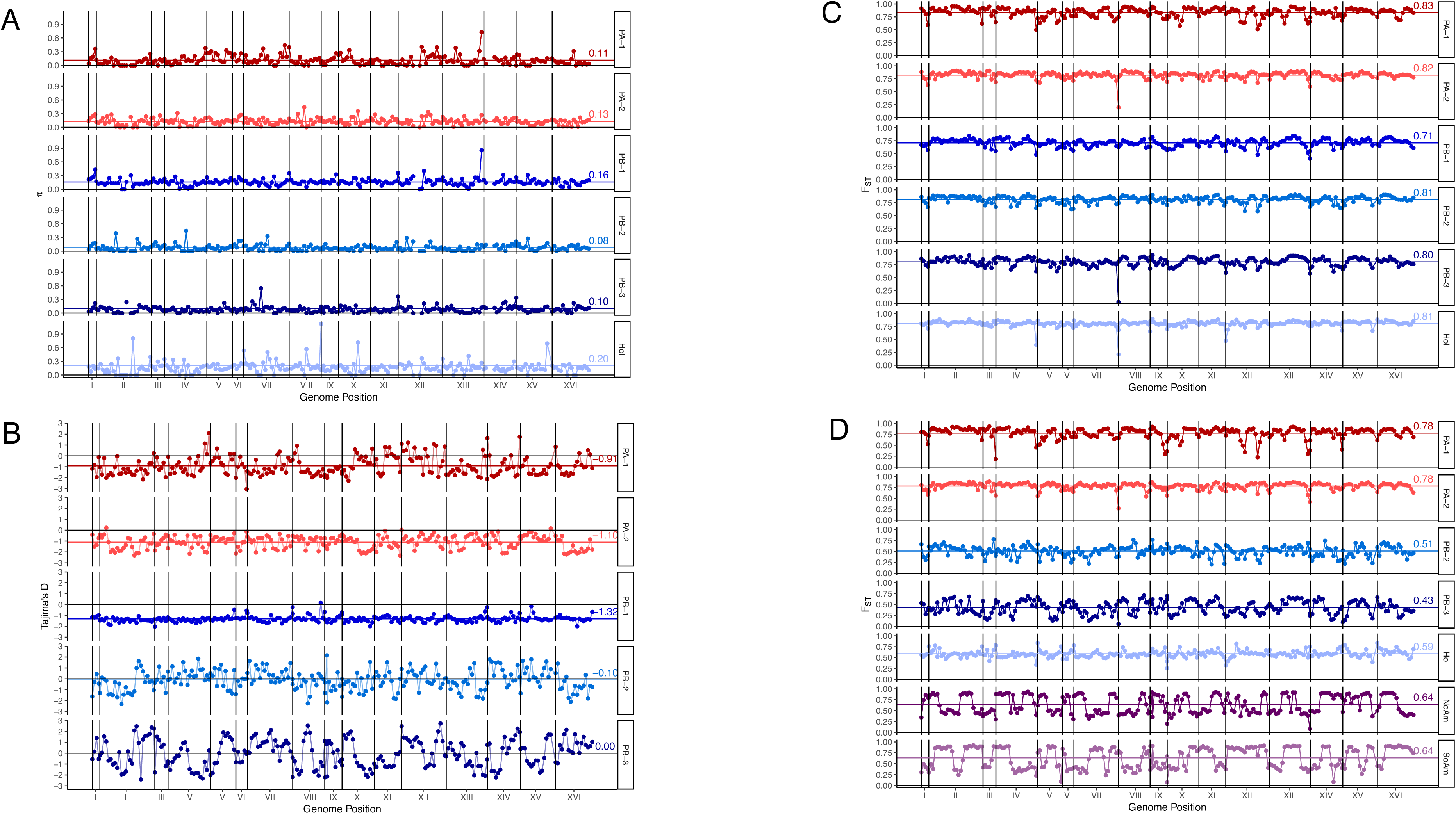
Additional population genomic statistics. (A) Mean pairwise nucleotide diversity (*π* * 100) for each subpopulation across the genome in 50-kbp windows. (B) Tajima’s D across the genome in 50-kbp windows for each subpopulation. (C) Mean F_ST_ in 50-kpb windows for each subpopulation compared to all subpopulations. (D) Pairwise F_ST_ for each subpopulation compared to PB-1.

**Supplementary Figure 5.**
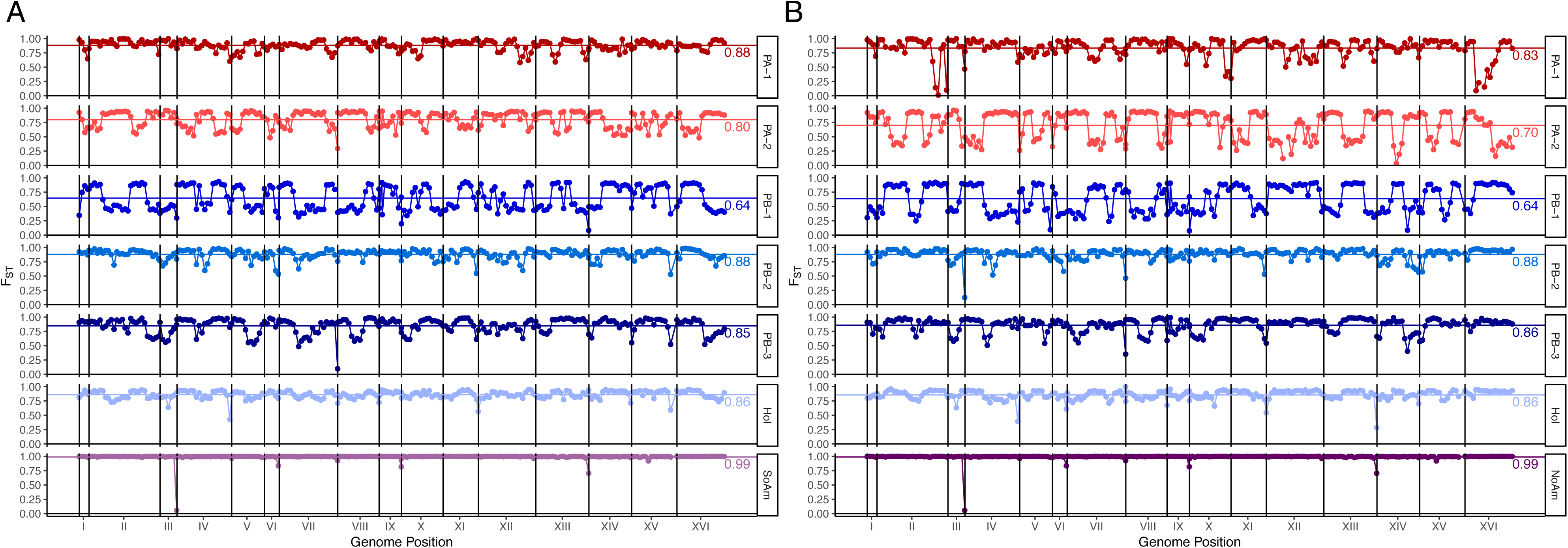
Pairwise F_ST_ plots for NoAm and SoAm compared to all other subpopulations. Pairwise F_ST_ for the NoAm lineage (A) or SoAm strain (B) compared to all other subpopulations.

**Supplementary Figure 6.**
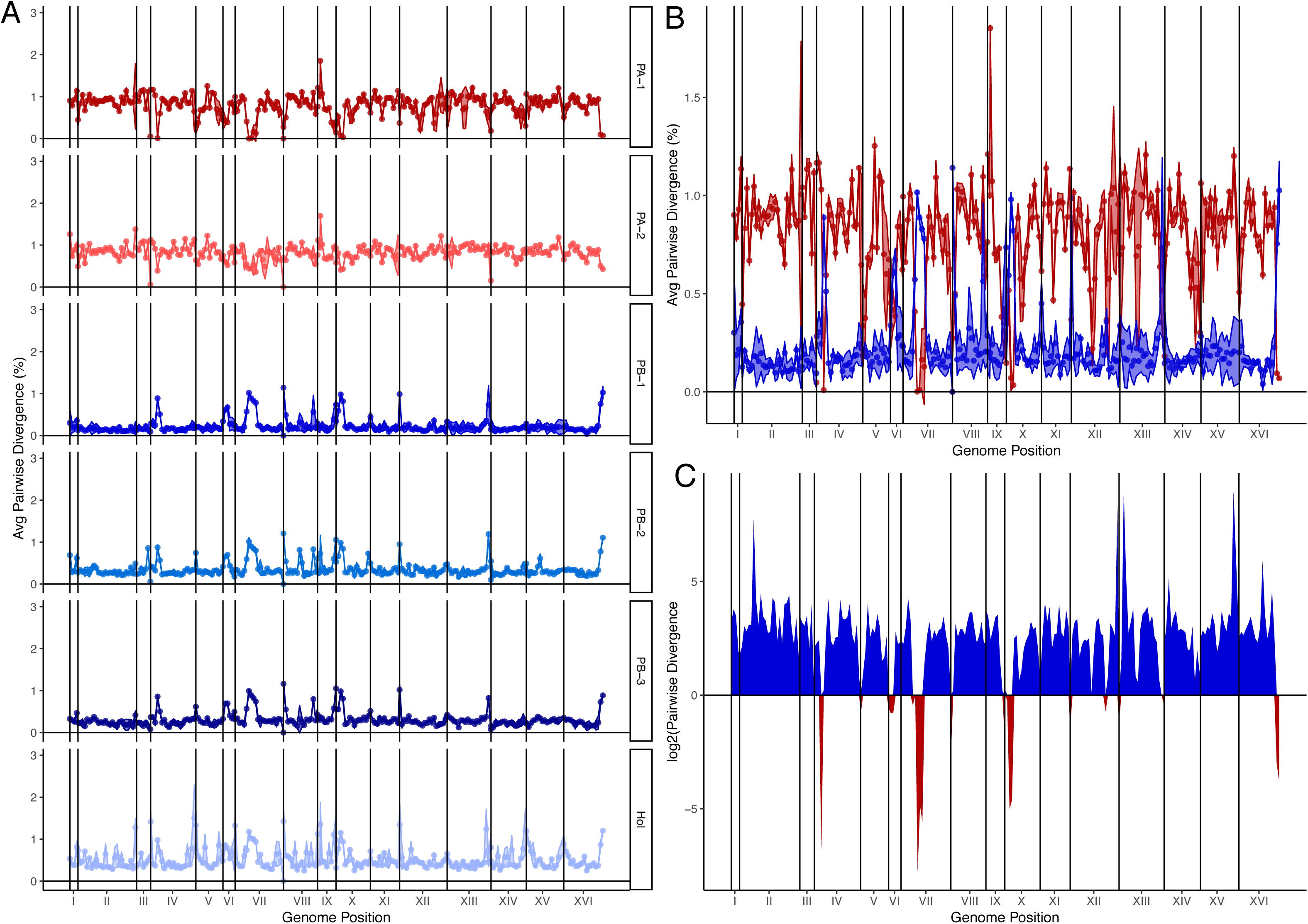
The taxonomic type strain has a mosaic genome. (A) Pairwise genetic divergence of the taxonomic type strain compared to each subpopulation. (B) Comparison of pairwise genetic divergence of the taxonomic type strain compared to PA-1 and PB-1. (C) log_2_ divergence plot (as in Figure 4) showing regions introgressed from PA-1 in the taxonomic type strain.

**Supplementary Figure 7.**
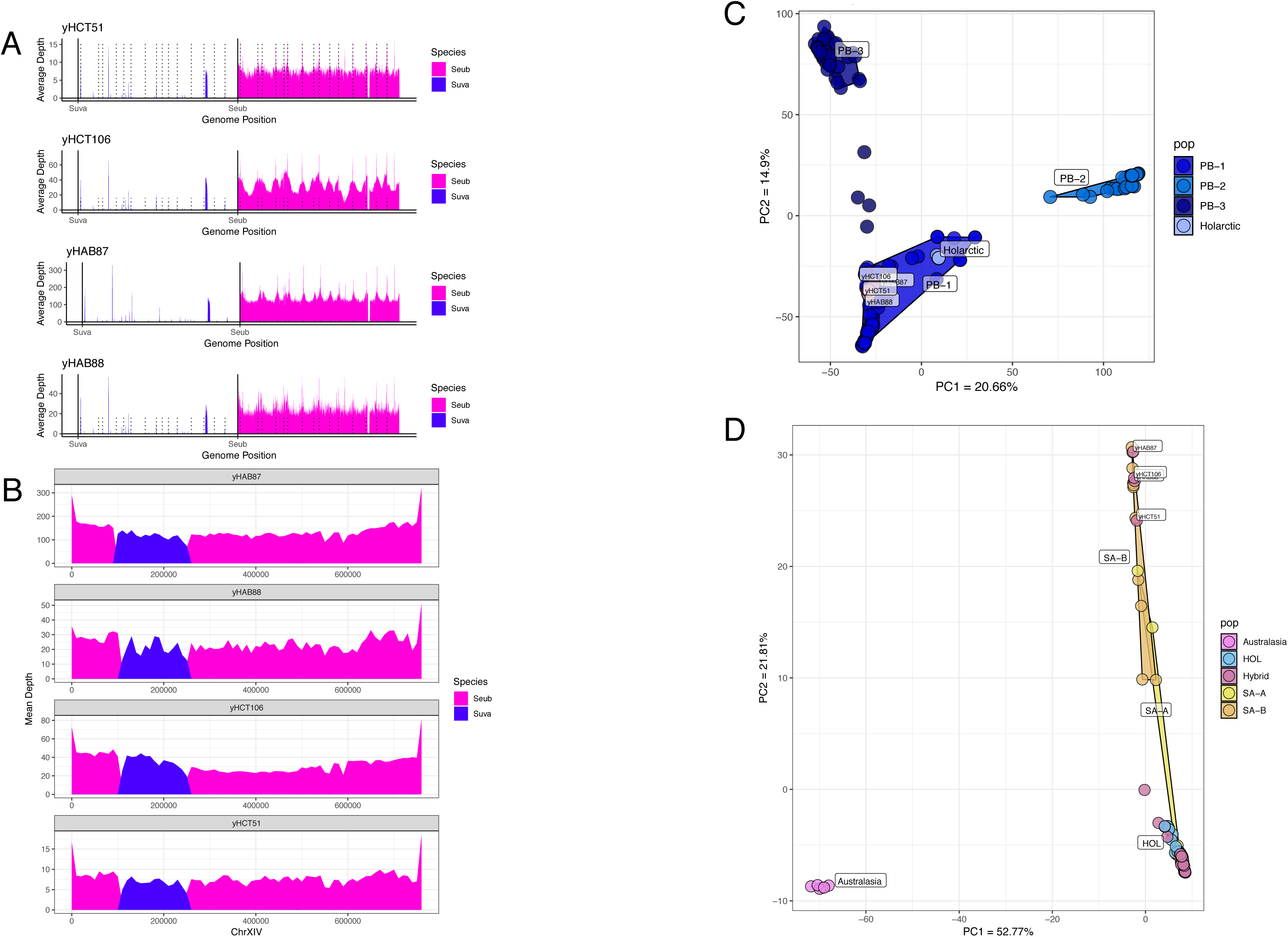
Four *S. eubayanus* strains with *S. uvarum* nuclear introgressions. (A) Depth of coverage plots of reads from four strains mapped to both the *S. uvarum* (Suva) and *S. eubayanus* (Seub) reference genomes. (B) Zoom-in of region on Chromosome XIV where these four strains have the same *S. uvarum* (purple) introgression into a *S. eubayanus* background. (C) A PCA plot shows that these four strains belong to the PB-1 subpopulation of *S. eubayanus*. (D) A PCA plot shows that the introgressed region from *S. uvarum* came from the South American SA-B subpopulation of *S. uvarum*.

**Supplementary Figure 8.**
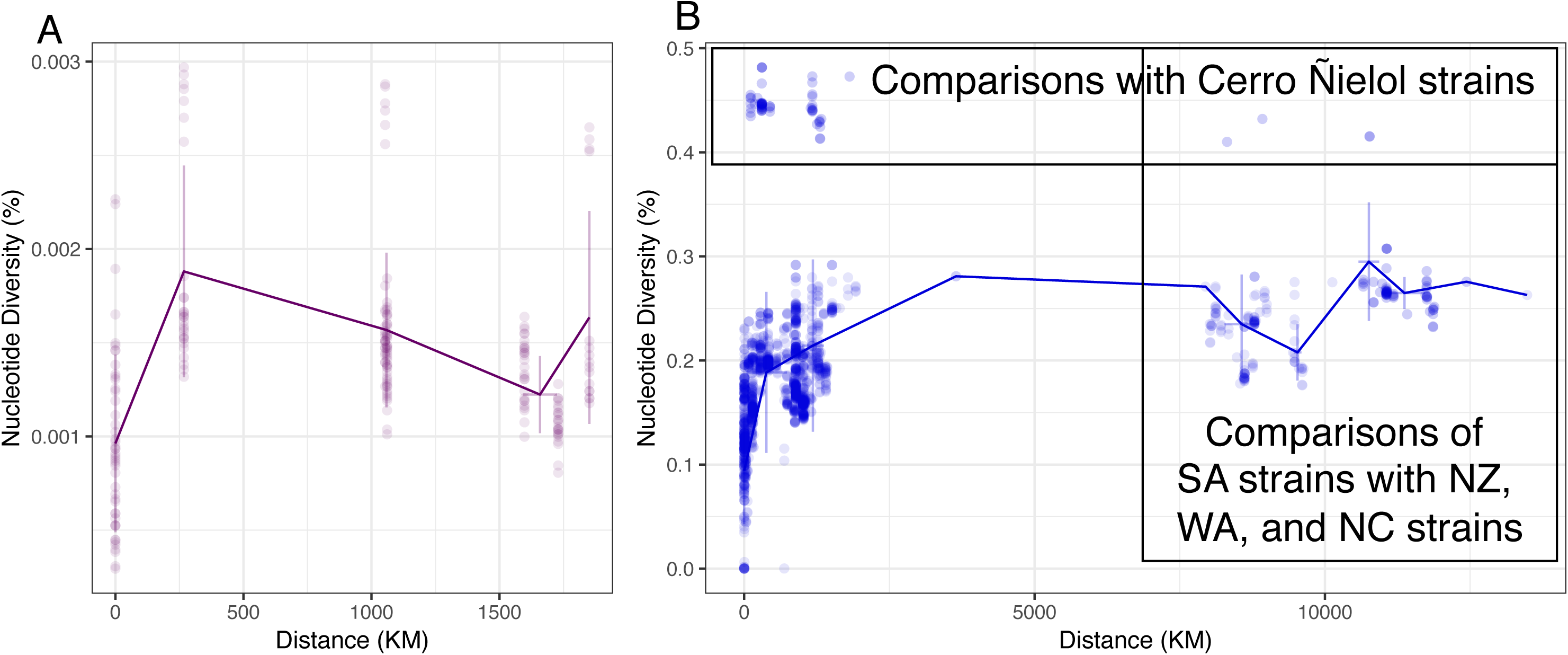
Isolation by distance plots for NoAm and PB-1. (A) Isolation by distance for all NoAm strains. The y-axis has been rescaled compared to Figure 5 for better visualization. (B) Isolation by distance for subpopulation PB-1. Comparisons with strains from Cerro Ñielol are labeled. All comparisons of South American strains with non-South American strains are on the right side.

**Supplementary Figure 9.**
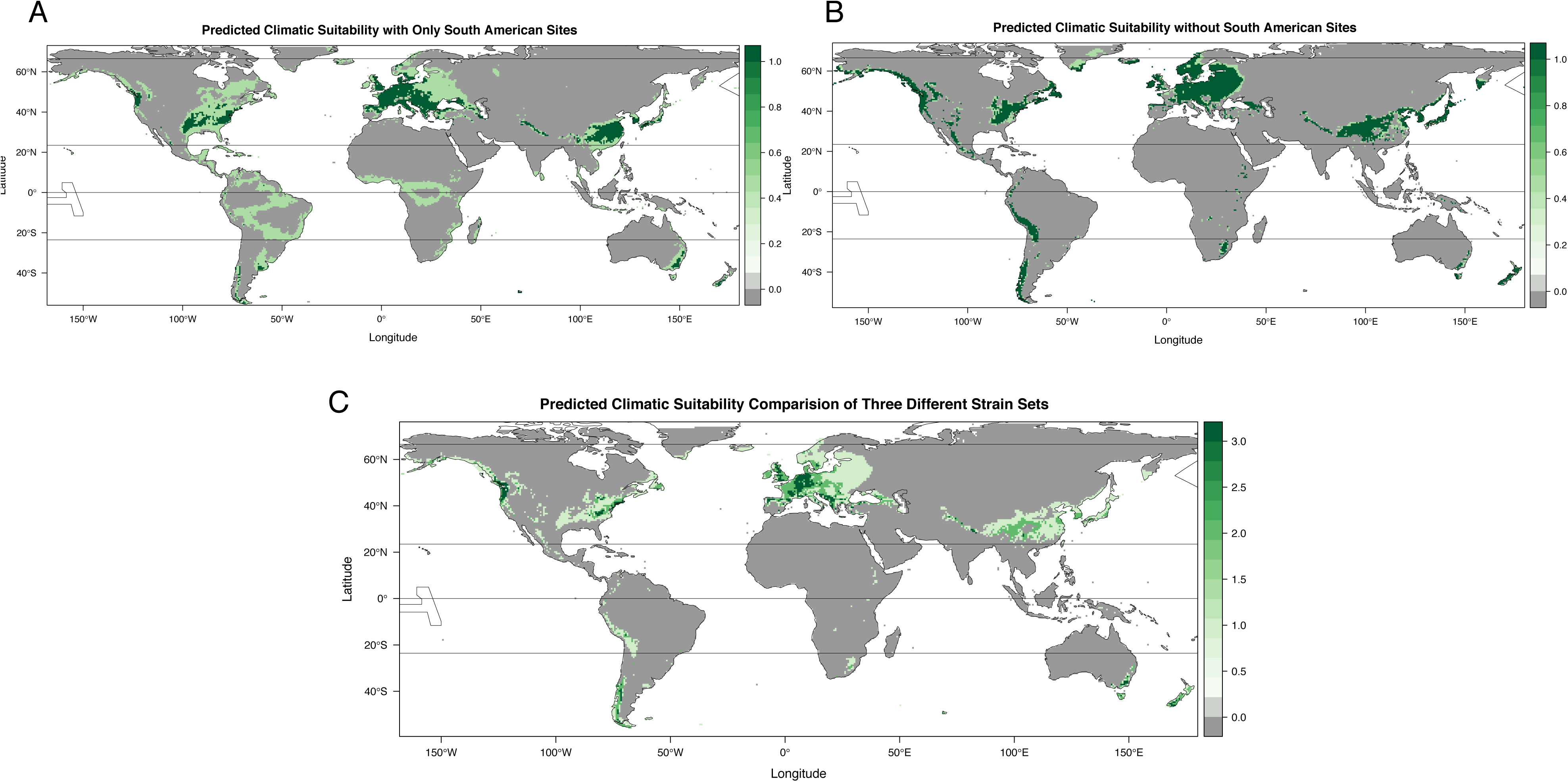
Additional Wallace climatic models. (A) Model built using only South American isolation locations. (B) Model built using only non-South American sites. (C) Comparison of models based on all known *S. eubayanus* collection sites, only South American, or only non-South American sites. Where the models agree is in dark green, where two models agree is in medium green, and where one model predicts suitability is in light green.

Supplementary Table 1. Collection information for all strains whose genomes were sequenced or analyzed in this study.

Supplementary Table 2. Average triplicate growth rates for various temperatures and carbon sources. Note that this spreadsheet has multiple sheets.

Supplementary Table 3. K=6 output of FastSTRUCTURE.

Supplementary Table 4. Mantel test results.

Supplementary Table 5. Input and output for Wallace climatic modeling.

